# Cannabidiol reverses fentanyl-induced addiction and modulates neuroinflammation

**DOI:** 10.1101/2024.07.20.604441

**Authors:** Bidhan Bhandari, Henrique Izumi Shimaoka Chagas, Sahar Emami Naeini, Pablo Shimaoka Chagas, Hannah M Rogers, Jules Gouron, Aruba Khan, Lívia Maria Maciel, Mohammad Seyyedi, Neil J MacKinnon, Hesam Khodadadi, Évila Lopes Salles, David C Hess, John C Morgan, Jack C Yu, Lei P Wang, Babak Baban

**Author notes:** These authors contributed equally to this work and should be considered as co-first authors (Bidhan Bhandari, Henrique Izumi Shimaoka Chagas). **Corresponding author:** Babak Baban, Ph.D., MPH, MBA, FAHA., -DCG Center for Excellence in Research, Scholarship and Innovation (CERSI) Augusta University, Augusta, GA, USA., -Department of Oral Biology and Diagnostic Sciences, DCG, Augusta University, Augusta, GA, USA., -Department of Neurology, Medical College of Georgia, Augusta University, Augusta, GA, USA., -Department of Surgery, Medical College of Georgia, Augusta University, Augusta, GA, USA.

## Abstract

**Introduction:** Fentanyl and non-pharmaceutical fentanyl use have been the leading causes of opioid-induced death worldwide. Being 50 times stronger than heroin and 100 times stronger than morphine, fentanyl is a potent opioid with overdoses causing over 250,000 deaths since 2018 in the US alone. The treatment of fentanyl addiction is a complex process and a clinical challenge. There is a dire need to find other innovative and alternative modalities in the fight against fentanyl crisis.

Increasing evidence suggests a correlation between neuroinflammation and symptoms of drug abuse, opening up the possibility of immunoregulatory agents as therapy for fentanyl addiction as well as a other opioid-induced addiction.

Cannabidiol (CBD) is a non-opioid, relatively safe, non-psychoactive phyto-cannabinoid produced by cannabis plants. Importantly, recent reports have documented benefits of CBD in the treatment and management of complications related to opioid withdrawal.

We investigated if inhaled CBD could reverse the fentanyl addiction and whether the CBD treatment could ameliorate the addiction symptoms by regulating neuroinflammatory signals and re-establishing the homeostasis in CNS.

**Method:** We used a fentanyl-induced conditioned place preference (CPP) model in mouse to test whether inhaled CBD could reverse the fentanyl addiction and ameliorate the adversarial symptoms. By employing a combination of flow cytometry as well as behavioral tests, we further assessed the impact of fentanyl addiction on cells and neuroinflammatory signals in CNS and we measured the effects of CBD in the treatment of addiction symptoms and inflammatory signals.

**Results:** Our findings suggest that CBD inhalation could be used effectively in the treatment of fentanyl addiction. CBD mitigated the excessive fentanyl-induced neuroinflammatory responses and decreased cellular stress and senescence.

**Conclusion:** inhaled CBD could alleviate the fentanyl addiction and regulate neuroinflammatory responses. This novel approach is non-invasive, accessible, effective, and warrants further, translational and research.

## Introduction

The opioid epidemic began as a healthcare system problem over 20 years ago in western countries due to overuse of synthetic opioids. The transformation of the opioid epidemic to a pandemic status is one of the most challenging crises of the current century with devastating effects, high mortalities, and heavy economic burdens on the healthcare system and communities (1-3).

In the last few years, fentanyl and non-pharmaceutical fentanyl use have been the leading causes of opioid-induced death worldwide (4,5). Fentanyl, a synthetic opioid, was initially invented by Paul Jansson in the 1960s for anesthetic and analgesic purposes (6). Being 50 times stronger than heroin and 100 times stronger than morphine, fentanyl is a potent opioid from which overdose has been responsible for over 250,000 deaths in the United States (US) since 2018. Fentanyl and its chemically similar analogues have been associated with a spike in deaths from opioid overdose, claiming 200 lives daily in 2022 in the US alone (7,8).

Treatments and management of fentanyl dependency, overdose, and withdrawal are challenging due to the complexity, personnel and length of therapy required, and cost. (9-11). Generally, opioid withdrawal, both acutely and in protracted manners, causes severe symptoms and, on occasions, irreversible clinical consequences including diaphoresis, tachycardia, extreme nervousness, anxiety, pain and insomnia, and digestive and metabolic disorders. A single direct detoxification approach has neither the expected effectiveness nor safety and could be accompanied by side effects and distress symptoms associated with withdrawal process (12). Medication-assisted treatment (MAT), in combination with behavioral therapies and counseling, is a more holistic therapeutic approach in the treatment of fentanyl overdosing and dependency. Despite the central role of MAT in the management of opioid addiction and prevention of mortality due to overdose, there remains major shortcomings associated with MAT. One of the main limitations is that many of the drugs used for it are themselves opiates, which may cause a new addiction, overdose, and relapse, and ultimately lead to treatment failure. Other problems include side effects, diversion from treating the main underlying causes of addiction/overdosing, and mis/abusing the medications by selling them as recreational substances (9-14). Therefore, there is a dire need to find alternative and innovative ways to combat the fentanyl crisis that are safer, non-invasive, non-opioid, cost effective, accessible, economic, highly efficacious, and with minimal to zero side effects.

Homeostasis is state of balance in living organisms by which the integrity and functional features of vital biological systems are maintained (15-17). The bidirectional communication between Central Nervous System (CNS) and Immune System (IS), neuroinflammation, plays a major role in dynamic adjustment of internal microenvironments to preserve the homeostasis and tissue stability (15-23). Within CNS, mounting evidence suggests a major role for microglia as a central hub for cellular manipulation and regulation of neuroinflammation (24-26). Importantly, there are several neurotransmitters and gliotransmitter receptors expressed by microglia which may contribute to their activation during addiction (27). Given the potential correlation between inflammatory responses and drugs of abuse (28), therefore, it is plausible to suggest that activation and polarization of microglia could be used as a therapeutic target in the containment of neuroinflammation and re-establishment of homeostasis in the CNS.

Further, excessive neuroinflammatory responses could affect the homeostasis negatively through metabolic irregularities and accelerated cellular senescence (29-32). Immoderate and exaggerated cellular senescence could drive the pathologic processes leading to vascular and neurodegenerative diseases (33-35). Therefore, exploring novel alternative and effective immunomodulatory strategies to sustain the homeostasis is crucial in the battle against drug abuse and addiction.

Cannabidiol (CBD) is a non-opioid, relatively safe, non-psychoactive phyto-cannabinoid produced by cannabis plants. There are numerous studies including recent findings from our group suggesting beneficial effects of CBD in several diseases through immunomodulation (36-38). There is an evolving notion that CBD could be used in the treatment and management of complications related to the opioid withdrawal process (39). Furthermore, several studies have reported potential use of CBD in the treatment of fentanyl addiction alone or as a supplement to existing therapeutic regimens (40,41). Given regulatory features of CBD, it is plausible to propose using such modulatory functions of CBD as potential alternative and/or adjunctive agents in the treatment and management of the opioid epidemic.

In this study, we tested whether CBD could reverse the fentanyl addiction. We further evaluated if CBD could modulate fentanyl-induced neuroinflammation, ameliorating adverse effects and improve outcomes. To the best of our knowledge, this current study is the first to use inhaled CBD to treat fentanyl induced addictive disorder. Importantly, the findings presented here pose neuroinflammation and its components as immunotherapeutic targets in the treatment of fentanyl addiction, transforming traditional modalities of opioid addiction by using an immunologic approach.

## Materials and methods

### Animals

Male C57BL/6 mice purchased from Jackson Laboratories USA were used in these experiments. Mice were 25-30 g in weight, aged 10-12 weeks at the beginning of the experiment, total of 30 mice from 3 independent cohorts. They were housed in a temperature-controlled room (22 ± 2°C) maintained on a 12 h light/dark cycle. The experiment began after a 1-week adjustment period. All animals were handled according to the National Institute of Health (NIH) guide for the Care and Use of Laboratory Animals. All experiments were conducted under the approval of the Augusta University Animal Care and Use Committee (Protocol # 2011-0062).

### Procedures and apparatus

#### a) Drugs and treatment

Fentanyl (1 mg/mL, Cerilliant^®^, USA), and inhaled broad spectrum CBD (ApelinDx^®^, Thriftmaster Holding group, TX, USA) (10 mg/mouse/day, 4 actuations) were used. Fentanyl was prepared by diluting 0.1mL of the original solution in 99.9mL of an isotonic solution of phosphate-buffered saline (PBS) (pH ∼ 7.4), resulting in a final fentanyl concentration of 1 μg/mL. Mice received by intraperitoneal injections 4μg/kg/day/mouse during drug adminstration days. For the absence of drug pairing during Conditioned Place Preference (CPP) conditioning, mice received PBS alone. The control group for addiction treatment inhaled purified air (generously provided by Thriftmaster Holding group specifically for this study), which was administered through inhalation the same way the CBD inhaler deployed.

#### b) Conditioned Place Preference

CPP is a well-established method widely used to investigate the reinforcing effects of drugs and drug craving. CPP is performed by using a two or three chamber apparatus, in which is possible to condition the subjects to associate a specific drug and the absence of said drug with the respective chambers of the apparatus, thus evaluating the abuse potential of substances and rewarding effects of drugs in mice through visual and tactile differences.

For this study, we used a CPP chamber meticulously designed and made by our collaborators at Georgia Institute of Cannabis Research of Medicinal Cannabis of Georgia LLC (Augusta, GA USA) (Figure 1). Our CPP chamber consisted of two equal-sized chambers (20 cm length x 15 cm width x 10 cm height) separated by a removable median partition (6 cm length x 15 cm width x 10 cm height). The chambers had a combination of floors (rough and smooth surface) and visual cues on the inside of the walls (black and white stripes in equal proportion, with horizontal or vertical patterns), and the median partition followed the same patterns. The temperature and light intensity were identical in both chambers. The mice were allowed to freely move between a compartment in which they were conditioned with either floors and/or cues.

**Figure 1.**
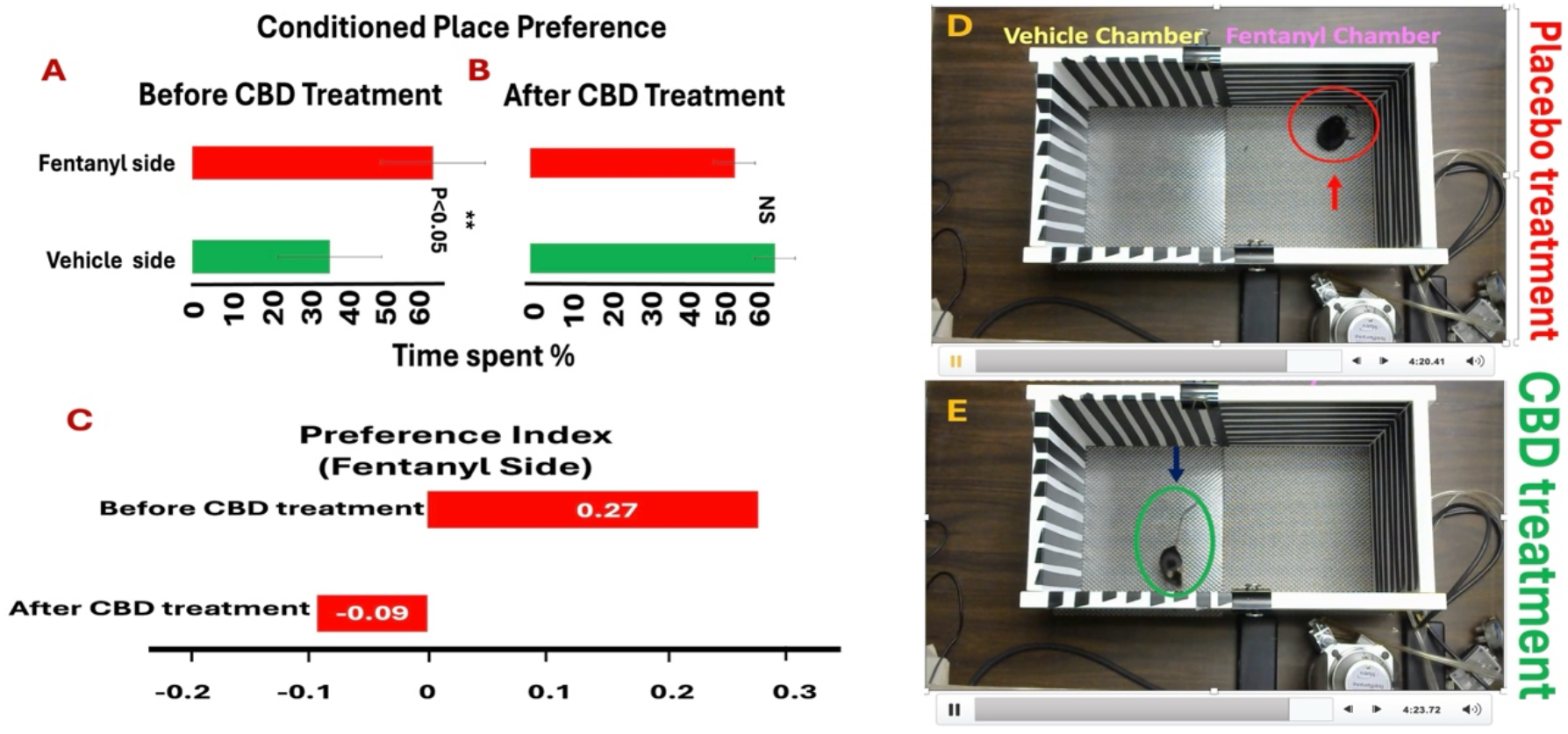
CBD inhalation reversed fentanyl addiction and palliated fentanyl adversarial symptoms. A) Fentanyl-induced conditioned place preference (CPP) was treated effectively by CBD inhalation **(B)**, reduced fentanyl addiction compared to counterparts treated with vehicle only (*p*<0.05). **C)** CBD inhalation reduced the PI significantly from +0.27 to −0.09 indicating a preference for the vehicle (non-fentanyl) side. **D-E)** Inhaled CBD not only reduced the fentanyl addiction, but also improved the side effects and symptoms of fentanyl effects such as abnormal body’s curvature stance and S shape of tail.

We used a modified unbiased three-stage CPP protocol. Briefly, at the first stage, all mice were preconditioned by being given free access to both chambers with no median partition for 10 minutes (day 0). The time spent in each compartment was recorded in order to determine the Initial Preference of each mouse. Fentanyl was associated with the least preferred chamber, whilst the absence of drug (PBS, vehicle) was associated with the most preferred chamber (Initial Preference) so we could avoid bias of each individual initial preference for one or the other chambers. The second stage of conditioning was for eight days (days 1-8). Experiments were conducted always starting at the same time of the day, and the mice were weighed daily so the dose of fentanyl could be given accordingly. On odd days of conditioning, all animals were intraperitoneally (i.p.) injected with our fentanyl solution and placed immediately in the associated drug chamber while, on even days, all animals received i.p. injection of PBS (0.1mL) and placed immediately in the vehicle chamber. After each injection, subjects were left in the chamber for 10 minutes with no access to other chamber using the median partition. On day 9, at the third stage of CPP, postconditioning, all animals were evaluated for CPP verification by removing the median and allowing animals to explore both chambers freely for 10 minutes. The time spent in each chamber during the preconditioning and postconditioning was measured and recorded. The CPP was verified and scored by comparing and measuring the time spent on each side (fentanyl or PBS chamber) during preconditioning minus postconditioning and was determined when there was a change in Final Preference regarding Initial Preference.

#### c) Preference Index (PI)

PI is a well-established method widely used to investigate the reinforcing preference index measured as the proportion of time spent on the side of the similar stimulus (42). It was calculated as the proportion of time the animals spent in the fentanyl side divided by the total amount of time spent in the fentanyl and vehicle sides together. A preference index significantly larger than 0.5 corresponds to a preference for the similar stimulus, and a value lower than 0.5 corresponds to a preference for the dissimilar stimulus. The higher the value, the stronger is the preference.

#### d Behavorial tests

##### d.1) Elevated Plus Maze (EPM)

the EPM is a well-established method to assess the anxiety/anti-anxiety effects of drugs. There are several variations of the experiments, but all of them aim to evaluate the time that subjects spent in conditions of exposure to threats, here represented by two opposite open arms, relating it with the amount of time spent in safer conditions, represented by two opposite closed arms, in a plus-shaped apparatus, elevated 1 meter from the floor. Each arm was 30 cm X 5 cm, and joined at a 5 × 5 cm central platform. The walls on the closed arms are 17 cm in height, closing all three corners of each closed arm. Open arms (OA) do not have walls. While exposing themselves in the open arms, the animals show an anti-anxiety behavior, in which they allow themselves to explore more the apparatus and the ambient. On the other hand, the more time spent in the closed arms the more anxiety behavior they display, aiming to stay safely close to the walls. Also, the higher the number of times that they cross from one arm to another the more anti-anxiety is their behavior.

For the EPM experiment a group of ten mice with CPP confirmed was evaluated. The apparatus was placed in the same room where the CPP experiments were conducted. Each animal was placed at the central space, all of them facing the same closed arm, and then recorded for 10 minutes.

#### e) Straub tail reaction

Injection of opiates, specifically morphine, induces Straub tail response in mice which indicates the drug-induced transient spasticity. Straub reaction is to assess the potency of centrally acting muscle relaxants. Here, in this study, we tested whether fentanyl could elicit such reaction as a measure for drug effects and potency in mice. After fentanyl injection, animals were directly observed, and tail responses were recorded immediately (within 10 minutes of observation). A modified intensity score scale was used to grade the Straub tail reaction as described previously (43). Accordingly, the grading system was from 0 to 4, 0 for no response (<30°); 1 for 30–45°above the horizontal table; 2 for 46–90°; 3 for more than 90°; and 4 for S-shaped curve of the tail.

#### f) Analytical Flow cytometry and immunophenotyping

Fresh brain tissues were processed and analyzed using flow cytometry as described previously with slight modifications (44). Briefly, whole brain was carefully harvested. Skin and muscles were removed, and the skull was carefully cut using angled scissors, avoiding damage to underlying brain tissue. Collected tissues were dissociated with the aid of a wide-tip plastic plunger and filtered through a 70 μm nylon cell strainer prior to further analysis. All single cell suspension samples were subjected to phenotypic and functional analysis of microglia (CD45^lo^CD11b^+^CD68^+^Iba1^+^IL-1b^+/-^ IL6^+/-^) and sub-populations of leukocytes including macrophages and their two subtypes (M1 & M2)(CD45^hi^CD11b^+^CD68^+^LY6C^+^LY6G^-^CD206^+/-^TNFα^+/-^IL10^+/-^, as well as lymphoid T cells (CD3^+^CD4^+/-^CD8^+/-^FOXP3^+/-^INF□^+/-^IL10^+/-^). Further analysis was performed by measuring cellular stress and senescence markers including T-cell Intracellular Antigen-1 related protein, TIAR-1 (Santa Cruz animal health USA) and P53 (Bioss, USA). All other antibodies were purchased from Biolegend. Cells were then run through a NovoCyte Quanteun flow cytometer. Cells were gated based on forward and side scatter properties and on marker combinations to select viable cells of interest. Single stains were used to set compensation, and isotype controls were used to determine the level of nonspecific binding. Analysis was performed using FlowJo (version 11.0) analytical software. Cells expressing a specific marker were reported as a percentage of the number of gated events. A population was considered positive for a specific marker if the population exceeded a 2% isotypic control threshold. The FlowJo plugin for the algorithm “uniform manifold approximation and projection” (UMAP) was used to perform and display the pattern distribution of double expression of cellular stress and senescence data sets.

#### g) Statistics

For statistical analysis, data were analyzed using Graphpad Prism 9. A one-way analysis of variance (ANOVA) followed by Kruskal-Wallis multiple comparisons test was performed to establish significance (*p* < 0.05) among all groups followed by Tukey’s test.

## Results

### Inhaled CBD enervated the rewarding effects of fentanyl addiction

As shown in figure 1A, CPP evaluation confirmed the phenotypic features of fentanyl addiction. All mice spent over 50% of their time in fentanyl side of CPP chamber. Such addiction to fentanyl was significantly reduced by treatment with inhaled CBD (Fig 1B), mice spent less than 30% in fentanyl side and the most at the vehicle side of the CPP chamber (\*\**p* < 0.05). Further, CBD inhalation reduced the PI significantly from +0.27 to −0.09 (Fig 1C). Additionally, fentanyl addiction induced specific phenotype and unusual behavioral characteristics including, not limited to, abnormal body’s curvature posture, and tail stance (S shape) with ataxia (Fig 1D). Such features of fentanyl addiction were significantly alleviated and improved by CBD treatment towards the normal position and deportment (Fig 1E).

### CBD inhalation reversed the fentanyl-induced Straub tail reaction and improved fentanyl associated stress symptoms

Addiction to fentanyl induced Straub response characterized by erected and inelastic tail across the back of the animal in an unusual S-shaped curve (Figs 1D and 2A). Treatment with CBD inhalation reversed such response compared to placebo treated group (Figs 1E and 2B). The chart (Fig 2C) is the graded demonstration of fentanyl-induced Straub effect comparing placebo treated group (graded between 2 and 4) compared to CBD treated mice (graded < 2) (*p*<0.01). Lower grading after CBD inhalation means that inhaled CBD reduced Straub effect’s spasticity which could be interpreted as a sign of lessened stress with improved brain and neurological conditions. Importantly, Inhaled CBD reduced fentanyl-induced anxiety. As displayed by bargraphs (Fig 2D), Elevated plus maze (EPM) assessment system showed that inhaled CBD treatment attenuated the fentanyl-induced anxiety. CBD treatment of mice with fentanyl addiction made them to enter the open areas during EPM test more than the addicted mice treated with placebo, an indicator of less anxiety compared to their counterparts received placebo. Although the difference was not statistically significant, however as shown, the tendency was highly and readily noticeable. The lack of statistical meaningful difference might have been because the low number of subjects (*P*> 0.05).

**Figure 2.**
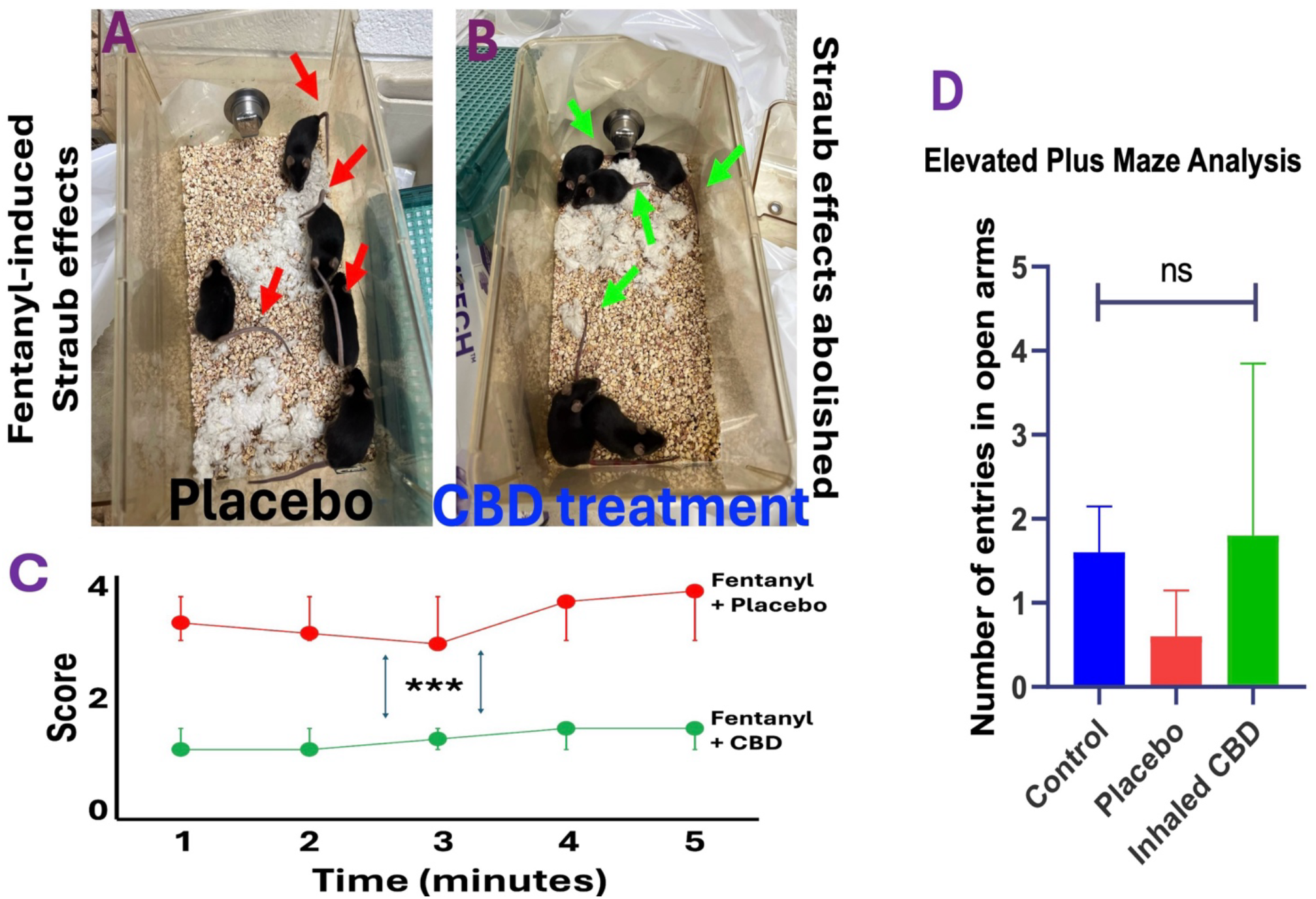
CBD inhalation assuaged fentanyl-induced Straub effects and improved adversarial symptoms including stress and anxiety. **A)** Fentanyl induced Straub effects featured by tail erection and inelasticity. **B)** Inhaled CBD was able to appease the Straub tail erection towards a normalized tail stance. **C)** Chart displays the grading of Straub tail effect induced by fentanyl. CBD inhalation was able to abolish the Straub effect, indicator of lessened stress (*p*<0.01). **D)** Using elevated plus maze, the level of anxiety induced by fentanyl addiction was measured. As shown in the chart, CBD inhalation was able to reduce the anxiety compared to the placebo treated group. Despite the obvious decreasing trend in CBD treated animals compared to placebo group, the difference was not statistically significant (*p* > 0.05), which may be due to the lack of power and limited number of subjects.

### Inhaled CBD reversed fentanyl-induced neuroinflammation

Flow cytometry analysis showed that fentanyl addiction altered the immune profile by increasing neuroinflammatory signaling. Single cell suspension from the whole brain tissues was analyzed to identify and measure microglia versus infiltrating macrophages and T cells based on their surface markers and intensity of CD45 expression (Fig 3). As shown in figure 3A, microglia were identified initially as CD45^lo^CD11b^+^ and further analyzed based on their expression of Iba1^+^ specific to microglia along with co-expression of IL-1b and/or IL-6 (figure 3B). As demonstrated, neuroinflammatory signaling including IL-6/IL-1 expression by microglia was reduced significantly by CBD treatment compared to placebo. Additionally, the flow cytometric UMAP analysis confirmed that CBD treatment not only reduced proinflammatory cytokines in fentanyl addicted mice, but also, altered the whole distribution pattern of neuroinflammatory signaling (Fig 3C). The quantified expression of IL-6/IL-1β by microglia is shown in figure 3D demonstrating significant difference between cytokine expression in CBD treated mice compared to placebo group (*P*<0.001). Further flow cytometry analysis showed CBD treatment shifted the profile of infiltrating macrophages (MQs) from M1 type MQs to M2, reducing the M1 inflammatory responses (Fig 3E). In addition, CBD inhalation was able to increase regulatory T cells, an indicator of anti-inflammatory function of CBD during fentanyl addiction (Fig 3F). Altogether, CBD inhalation showed anti-inflammatory effects during fentanyl addiction evidenced by reduction of neuroinflammatory signaling including lower microglia-induced pro-inflammatory cytokines, increased infiltrating M2 MQs and higher regulatory T cells.

**Figure 3.**
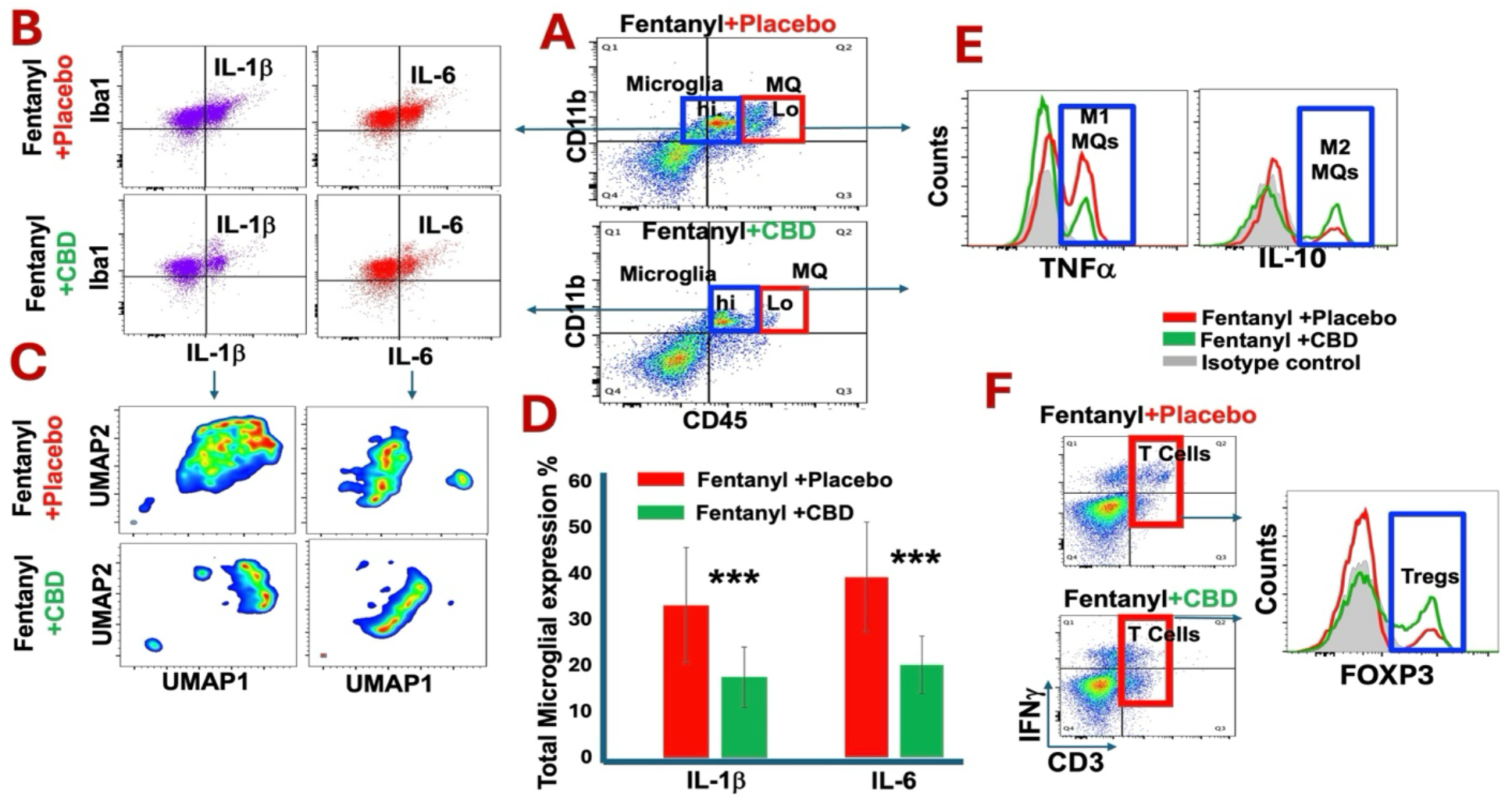
CBD inhalation downregulated fentanyl-induced neuroinflammation. An assessment of neuroinflammatory signals including microglia, infiltrating macrophages and T lymphocytes was performed using flow cytometry analysis**: A)** CBD inhalation could enhance counter inflammatory functions of both microglia (CD45^lo^CD11b^+^Iba1^+^) and infiltrating macrophages (CD45^hi^CD11b^+^ IL-10^+^) with higher level of IL-10 expression compared to placebo treated group. **B)** CBD inhalation reduced expression of pro-inflammatory cytokines, IL-1b and IL-6, by microglia compared to placebo group. **C)** The analysis of flow cytometry data using the FlowJo software plugin for the algorithm UMAP. The UMAP subtraction analysis revealed numerous spatial clusters with significant differences in IL-1b and IL-6 distribution between fentanyl addicted mice treated with placebo versus CBD treated mice. **D)** The chart with bargraphs demonstrates the quantified measures for IL-1b and IL-6 expression by microglia with significant reduction in CBD treated compared to placebo group (^***^*p*<0.01). **E)** CBD treatment reduced the type 1 infiltration macrophages (CD45^hi^CD11b^+^ TNFa^+^) while enhanced the type 2 with higher expression of anti-inflammatory IL-10 cytokine. **F)** CBD inhalation increased the frequency of regulatory T cells (Tregs: CD3^+^FOXP3^+^IL-10^+^IFNg^-/lo^) with higher expression of IL-10, promoting anti-inflammatory signals.

### Inhaled CBD mitigated fentanyl-induced cellular senescence and stress signaling

Flow cytometry analyses showed that fentanyl addiction increased cellular stress and senescence by enhancing expression of TIAR-1 and P53. CBD treatment reduced stress and senescence signaling (Fig 4A). Figure 4B demonstrates the analysis of flow cytometry data using the FlowJo software plugin for the algorithm UMAP. The UMAP subtraction analysis revealed numerous spatial clusters with significant distribution differences between fentanyl addicted mice treated with placebo versus CBD treated mice. The chart with bargraphs (Fig 4C) demonstrates the quantified measures for stress/senescence comparative levels in fentanyl addicted mice treated with placebo versus CBD treated mice with significant differences (^**^*P*<0.05).

## Discussion

The present study is the first report to show a potential role for CBD in the treatment of fentanyl addiction in an experimental model. Our findings demonstrated that inhaled CBD reduced the fentanyl addiction and improved the symptoms in animals with fentanyl addiction. Our data also indicated that inhalation of CBD was able to lessen Straub tail elevation effect. Straub phenomenon is an indicator of stress and spasticity, has been mainly described in morphine addiction (45,46). In this study, not only do we report that fentanyl addiction could induce Straub effect, but also our findings demonstrated for the first time that CBD inhalation was able to improve stress and complications related to Straub phenomenon. Such beneficial impact of CBD could potentially be extended to treat a broader range of opioid addictions and to improve the brain and motor nerves conditions.

**Figure 4.**
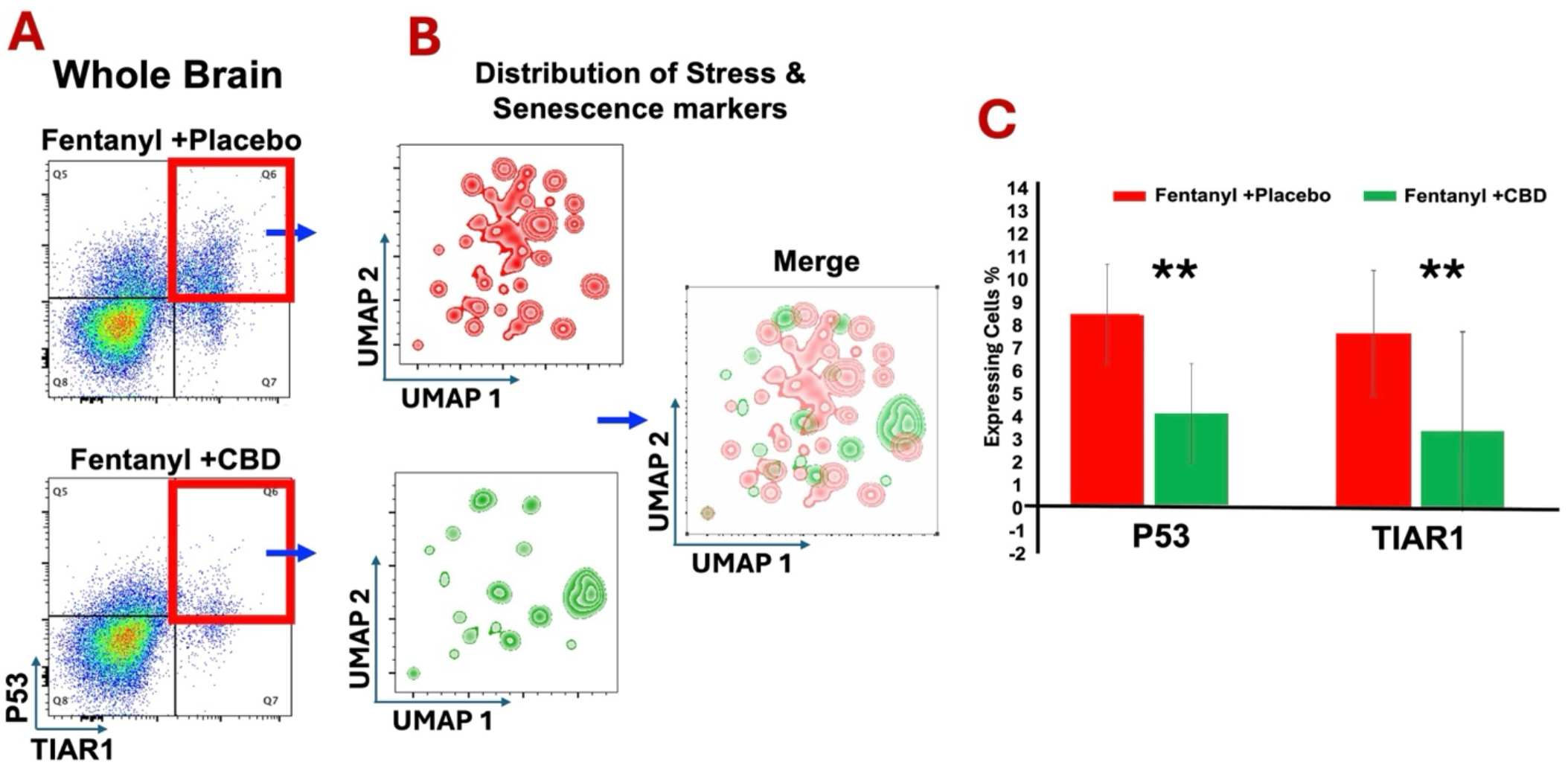
Fentanyl-induced cellular senescence and stress signaling were reduced by CBD inhalation. **A)** Flow cytometry analysis showed that CBD treatment reduced stress and senescence signaling (TIAR-1 and P53 respectively) compared to placebo treatment. **B)** The analysis of flow cytometry data using the FlowJo software plugin for the algorithm UMAP. The UMAP subtraction analysis revealed numerous spatial clusters with significant differences in the distribution of TIAR-1 and P53 signaling between fentanyl addicted mice treated with placebo versus CBD treated mice. Merged panel clarifies the visual differences in distribution. **C)** Bar graphs display the quantified measures (%) of total expression of TIAR-1 and P53, CBD inhalation reduced both signals significantly (*p*<0.05).

Several studies have used CBD in the treatment of addiction behavior and symptoms in both pre-clinical and clinical settings (47-50). The mechanisms by which CBD improves the conditions in addiction are unclear (51,52). However, some CBD’s properties including anxiolytic, anti-psychotic, antidepressant, and neuroprotective effects have been among factors that rationalize the use of CBD in the treatment of addiction. Additionally, the potential of CBD in manipulation of endocannabinoid system, internal signals and receptors (e.g., CB1, TRPV1, GPR55 etc.) contributes to the alleviation of addiction symptoms and treatment (53-56).

The findings reported in this current study suggest CBD inhalation is a potential effective treatment of fentanyl addiction with less challenges and side effects associated with established pharmacologic interventions. The exact mechanism of action by which inhaled CBD could reverse the adversarial effects of fentanyl addiction remain to be elucidated. However, importantly, for the first time, our study showed that CBD inhalation altered the tectonic plate of neuroinflammation plateau in CNS during the fentanyl addiction. There are significant number of studies indicating a link between drug-addiction and upregulation of pro-inflammatory responses in CNS (28,30). Such inflammatory microenvironment has been associated with addiction behavior, lack of coherent cognitive function, dysfunction of blood brain-barrier (BBB), neural pathogenesis, enhanced cellular senescence and immune imbalance. In our study, CBD inhalation was able to downregulate neuroinflammatory responses by affecting microglia, reducing their inflammatory cytokine production (IL-6/IL-1β) significantly and altering the distribution pattern of proinflammatory cytokines. Such shift was associated with reduced addiction to fentanyl, lower anxiety and importantly, attenuated cellular stress as well as senescence signaling. Further, our data showed that CBD inhalation could alter the polarization of infiltrating macrophages towards M2 type and enhanced Tregs within CNS. Microglia, peripheral monocytes (infiltrating macrophages) lymphocytes (T cells) are all immuno-inflammatory cells with variations in their origins and functions. Microglia are suggested to derive solely from marrow myeloid progenitors produced by yolk sac in embryonic period. Other CNS resident macrophage subsets arise later during embryonic development (57,58), while T Lymphocytes are originated from lymphoid lineage in bone marrow. Together, both microglia and the peripheral immune cells are increasingly recognized for their crucial roles in the development, homeostasis, and diseases of the central nervous system. Our data here in this study not only reveals the interplay between fentanyl addiction and neuroinflammation, but also underscores the significance of CBD as an immunotherapeutic target in regulating the neuroinflammation and in re-establishing the homeostasis in CNS affected by fentanyl abuse. Whether the fentanyl-induced impairment of neural circuit occurred prior to excessive neuroinflammatory responses or not, our data demonstrated that fentanyl addiction caused significantly increased inflammatory responses in CNS. Therefore, it is plausible that the beneficial effects of CBD shown in this study may well relate to the anti-inflammatory features of CBD. Additionally, inflammation is known to play a central role in cellular stress and senescence signaling (59-61). The immunoregulatory feature of CBD could explain the reduction of stress and cellular senescence in CBD treated mice. This warrants further research.

In conclusion, the fentanyl addiction is the major opioid epidemic worldwide. This is due to fentanyl’s high potency, ease of synthesis and mixing with other drugs and narcotics, and low cost. Despite relative efficacy, current pharmacologic modalities for addressing fentanyl addiction are associated with significant challenges and many shortcomings (4,9,11, 62-65). Neuroinflammation is a major contributing factor of fentanyl addiction and its symptoms (28,66). The novel findings in this study not only propose an immunologic-based association for fentanyl addiction, but also suggest that CBD is a relatively safe, affordable, and natural immunotherapeutic modality in the treatment of fentanyl addiction with minimal to zero side effects and highly favorable outcomes. The data reported in this study are at preliminary stages with extremely high potential impacts, and requires more extensive, translational research in this area.

## Abbreviations

CPP: Conditioned Place Preference
CBD: Cannabidiol
CNS: Central Nervous System
MAT: Medication-assisted treatment
EPM: Elevated Plus Maze
BBB: Blood Brain-Barrier;
TIAR-1: T-cell intracellular antigen 1-related (TIAR-1) protein.

## Author contributions

BB, LPW: Conceptualization, Data curation, Formal analysis, Funding acquisition, Investigation, Methodology, Project administration, Resources, Software, Supervision, Validation, Visualization, Writing – original draft, Writing – review & editing. BIB, SEN, HMR, JG, PSC, LMM: Data curation, Formal analysis, review & editing. HISC, ÉLS: Data curation, Formal analysis, Investigation, Methodology, Project administration, Software, Validation, Visualization, Writing – review & editing. MS, NJM, DH, JCY: Investigation, Writing – original draft, Writing – review & editing.

## Acknowledgement

Authors are thankful to ThriftMaster Holding Group for providing the inhalant CBD for this study. Authors also thank Medicinal Cannabis of Georgia for providing help in optimizing the CBD dosage.

## Data availability statement

The original contributions presented in the study are included in the article material, further inquiries can be directed to the corresponding authors.

## Ethical publication statement

We (authors) confirm that we have read the Journal’s position on issues involved in ethical publication and affirm that this report is consistent with those guidelines.

## Funding

This work was supported by institutional seed funding from the Dental College of Georgia at Augusta University.

## Declaration of Competing Interest

1-Lei Phillip Wang, Babak Baban, and Jack Yu are members of Medicinal Cannabis of Georgia with no financial interest. 2-All other authors declare no conflict of interest. 3-Thriftmaster Holding Group (THG) is the provider of CBD inhalers and has a licensing contract with Augusta University. 4-THG and its members had no role in study design, data collection and analysis, decision to publish, or preparation of the manuscript.

